# AnimalMotionViz: an interactive software tool for tracking and visualizing animal motion patterns using computer vision

**DOI:** 10.1101/2024.10.22.619671

**Authors:** Angelo L. De Castro, Jin Wang, Jessica G. Bonney-King, Gota Morota, Emily K. Miller-Cushon, Haipeng Yu

## Abstract

Monitoring the movement patterns of dairy cattle can provide important insight into space utilization or space occupancy in a barn. Although several precision livestock technologies have been developed to record dairy cattle movements, there is a lack of open source tools to track and visualize cattle movement patterns. Therefore, we developed an open-source computer vision software tool, AnimalMotionViz, that allows users to track and visualize dairy cattle movement patterns using a motion heatmap. The software comes with an easy-to-use web-based graphical user interface built with the Python Dash package. It implements a set of background subtraction algorithms in the OpenCV package to track animal motion patterns in real time. The software processes each frame of the input video and identifies the background and foreground using these algorithms. Foreground objects are then subtracted from the background across all frames and cumulatively overlaid on an empty mask image created with the first frame of the input video to visualize the intensity or frequency of motion across different regions. The user can generate motion heatmaps in an image and video, and also track specific regional motion with a custom mask. The software also returns the top three peak intensity locations, the total percentage of regions used, and the within-quadrant percentage of regions used. In four 5 min sample videos, quadrants with peak intensity of space use, as identified using the software, aligned with quadrants where calves spent the greatest duration of time, according to continuous recording of behavior from video. The motion heatmaps generated by AnimalMotionViz can be used to understand space utilization or space occupation by animals, as well as to assess how space allocation affects dairy cow movement. We conclude that the newly developed AnimalMotionViz is a user-friendly and efficient tool to support research developments in precision livestock farming towards enhancing cattle management practices and improving pen designs.

## Introduction

The welfare and productivity of intensively housed dairy cattle is highly dependent on aspects of the housing environment (reviewed by Bewley et al., 2017) and behavior is often interpreted as a key indicator of the welfare implications of housing management factors. For example, resting and feeding time are sensitive to stall (Krawczel et al., 2012) and feed bunk (DeVries et al., 2004) stocking density, respectively. Space allowance influences patterns of rest and heterogeneity of space use in dairy cows (Marin et al., 2024), although effects of more general space allowance remain less studied in adult cows (Thompson et al., 2020), possibly due to challenges in quantifying effects of management on broader use of pen space. To date, much research has focused on individual behavioral indicators while providing less insight into group-level movements or space use.

Improved understanding of dairy cow pen space utilization has potential applications related to both refinement of housing management and assessment of animal welfare. Specific use of space provided (e.g., preference for perimeter of the pen in calves; Morita et al., 1999; Marin et al., 2024) may provide insight into preference or aversion for pen areas or specific features to refine decisions related to pen design. Similarly, information related to group-level or temporal variability in pen space use allows characterization of grouping or bunching behavior in dairy cows, which may be related to aspects of pen or barn design (van Schaik et al., 2021). Grouping behavior may additionally be subject to thermal conditions (Chopra et al., 2024), reflecting a potential group-level indicator of response to heat or cold stress. Space use is also influenced by individual factors, including social dominance (e.g., proximity to food sources; Nakanishi et al., 1992) and pain (e.g., increased use of a shelter in group-housed dairy calves following disbudding; Gingerich et al., 2020).

Tracking space use or occupancy may provide novel and useful information for improving livestock management, yet measurement of this data poses a challenge given limitations in visually monitoring all animals. Precision livestock farming uses a variety of sensor technologies and data-driven analytics to monitor or phenotype animals at the individual level (Morota et al., 2018). Computer vision is one of the subfields of precision livestock farming that analyzes images or video data from cameras. A recent study successfully used computer vision to estimate pen space utilization by creating motion heatmaps in Holstein calves (Marin et al., 2024). However, the use of computer vision to monitor animal space utilization is limited in the literature due to the lack of publicly available software tools. Thus, our objective was to develop an interactive, open-source computer vision software tool for tracking and visualizing animal motion patterns. Previous studies have shown that the use of an interactive graphical user interface (GUI) is useful to reach the user without specific training in data science (Morota et al., 2021) or computer vision (Wang et al., 2024). Here, we describe an interactive GUI based on Python Dash (Hossain et al., 2019), document the statistical methods and computer vision algorithms implemented in the software, and present examples that compare the output of the software with observations of space use and locomotor behavior.

## Materials and methods

### Overview of software architecture

AnimalMotionViz was developed entirely in Python, taking advantage of its versatility and ease of use to create a user-friendly software application. The Dash framework was used to build a GUI, chosen for its ability to streamline the creation of interactive and responsive software applications. The GUI of the AnimalMotionViz application is divided into input and output sections. The input section includes video processing parameters where the user can upload a video, select a color map, and adjust various settings. We replaced the file upload components in Dash, which do not support files larger than 200 MB, with the open source dash-uploader (https://github.com/fohrloop/dash-uploader), which has no file size limitation other than the user’s available disk space. The user input collected in this section is then passed to a callback function for processing and parsing, with the results returned to the GUI. The output section displays the results in two tabs and a summary table. The heatmap image tab displays the final processed motion pattern image, with the top three peak intensity locations highlighted with symbols. The heatmap video tab displays the processed motion heatmap video using the Flask framework, which is capable of efficiently streaming and serving large videos in the GUI. The summary table uses multiple metrics to quantify the motion patterns relevant to the motion heatmap. This structured layout ensures a clear and efficient presentation of both input parameters and output results, providing a comprehensive analysis of animal motion. The overall software workflow is illustrated in Figure 1.

**Figure 1.**
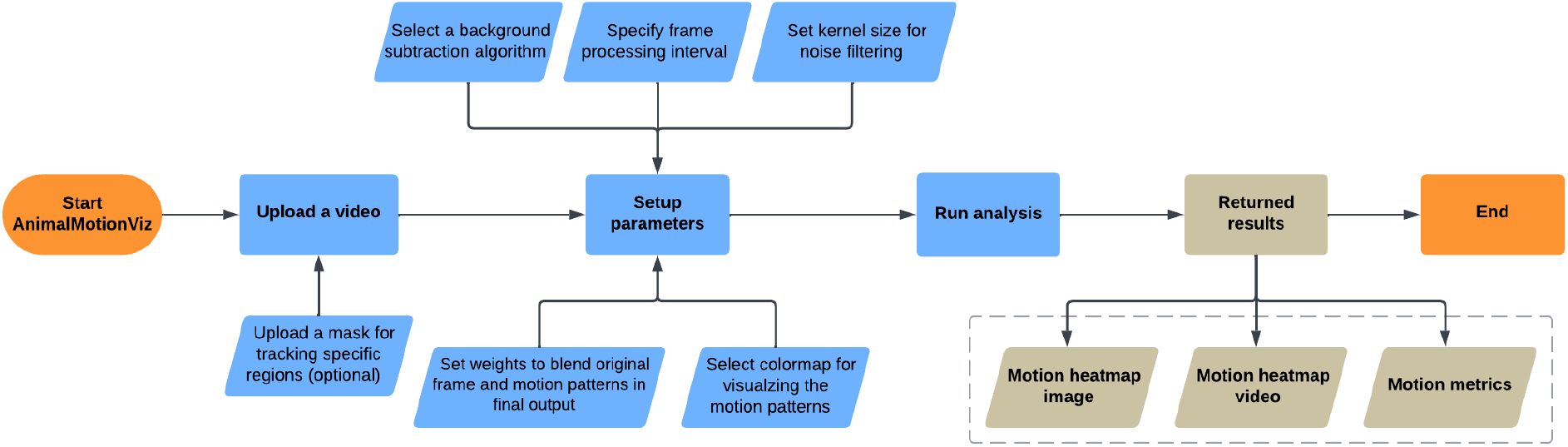
Overall workflow of AnimalMotionViz: The start and end points are highlighted by bright orange shapes, the user inputs are shown in blue shapes, and the returned results are represented by beige shapes.

### Implementation

The development and implementation of AnimalMotionViz relies on several libraries and frameworks to ensure the efficiency of the application in producing the intended results. A user-friendly GUI is built with Dash, a Python framework for building interactive software web applications. The design and layout of the web application were refined with the Dash Bootstrap Components, and the Dash AG Grid was used to create data tables in the Dash application. The dash-uploader component is adapted for efficiently uploading large video files. The main processing engines are OpenCV (Bradski, 2000) for handling image and video processing and NumPy (Harris et al., 2020) for numerical computation in Python. The base64 module was used to encode images for uploading, decode the uploaded image data for downstream analysis, and re-encode the final motion pattern image for embedding in HTML for display in the Dash web application. The imageio library (https://github.com/imageio/imageio) converts the processed images into a video file, while the Flask framework is used to stream and serve the converted video for display within the Dash web application.

### Inputs

The first step is to upload a video of interest using supported file formats, including mp4, avi, mov, wmv, mkv, and flv. If the user is only interested in tracking a sub-region of the video, they have the option of uploading a mask image created using annotation tools that allow a region of interest to be specified in the image, such as LabelMe (Russell et al., 2008) and Roboflow (Lin et al., 2022). While this step is optional, it is recommended as it allows the definition of specific areas of interest to be considered during video processing, thereby increasing the focus and relevance of the motion heatmap analysis.

The uploaded video is converted into images (i.e., frames) and then undergoes background subtraction to detect animal movement by separating the foreground animals from the static background. The user can choose from a number of algorithms implemented with OpenCV, including Improved Mixture of Gaussians (MOG2), K-Nearest Neighbors (KNN), Geometric Multi-Grid (GMG), CouNT (CNT), Local SVD Binary Pattern (LSBP), and Google Summer of Code (GSOC). These algorithms are known for their varying strengths in handling various environmental conditions, such as lighting changes and background variability. The default background subtraction algorithm implemented in the software is MOG2, an adaptive algorithm that models each pixel independently as a mixture of Gaussian distributions over time to separate the foreground and background, allowing it to handle dynamic backgrounds and illumination changes (Zivkovic, 2004; Zivkovic and Van Der Heijden, 2006). The KNN algorithm is a non-parametric algorithm, which stores the history of pixel values for each pixel location and uses the k-nearest neighbors approach to classify the background (Barnich and Van Droogenbroeck, 2010). The third algorithm, GMG, combines statistical background image estimation with Bayesian segmentation. It requires an initial learning period and is sensitive to noise, but can achieve high accuracy once the learning phase is complete (Godbehere et al., 2012). The CNT algorithm is a real-time counter-based algorithm that tracks how often each pixel location remains unchanged to classify the background and foreground, making it efficient for handling illumination changes (Dey and Kundu, 2013). The LSBP algorithm uses local binary patterns combined with singular value decomposition to perform background subtraction, making it effective for handling background noise and dynamic changes (Guo et al., 2016). The GSOC was introduced to make LBSP faster and more robust.

Additionally, the user can specify the image processing interval to process every *n*th image (the default interval is 1). The software also allows the user to specify the kernel size (ksize) to perform a morphological operation on the detected motion to reduce small noise. A small kernel removes small noise and preserves motion structure, while a larger kernel removes more noise but may also eliminate useful motion structure. The user can also adjust two weighting parameters and select a color map to visualize the detected cumulative motion patterns. The *α* and *β* weighting parameters are used to overlay the detected cumulative motion on top of the original frame to highlight the motion relative to the scene. This is achieved by combining the cumulative motion and the original frame with a weighted sum, where the *α* parameter is the weight of the original frame and *β* is the weight of the cumulative motion. To enhance the visualization of the cumulative motion, the user can select a color map from several color maps, including bone, ocean, pink, and hot, provided in the OpenCV library.

### Video processing

The software captures the uploaded video stored in local disk space and converts it to frames using OpenCV. If a mask file is provided, it is decoded from a base64-encoded string to binary data using the base64 module, and the decoded binary data is then converted to a Numpy array using the Numpy package for further analysis. The software then undergoes a user-selected background subtraction algorithm in OpenCV that distinguishes the foreground (i.e., moving animals) from the static background. Once the background is identified, the software processes frame by frame to detect motion and uses a binary image to represent the detected motion. The decoded and converted mask image (if available) is then applied to the motion image to restrict the motion detection to the specific region defined by the mask image using OpenCV. A morphological opening operation based on the user-specified ksize parameter is then used to filter out small noise to obtain a cleaner and more accurate motion. The binary mask representing the detected motion in a given region is then accumulated over the frame.

The recorded cumulative motion data is first sorted by pixel intensity, and the coordinates of the three pixels with the highest intensity values, representing the most active locations, are retrieved. This cumulative motion data is then used to calculate the total and within quadrant percentage of region used. The total percentage of region used is calculated as the total number of active pixels (i.e., pixel value > 0) divided by the total number of pixels in the given view. For the quadrant-specific calculation, the view is divided into four equal sections (quadrants) by drawing a horizontal and a vertical line that intersect in the center of the view. Quadrants 1 through 4 correspond to the upper right, upper left, lower left, and lower right regions, respectively. The percentage of area used is then calculated for each quadrant using the same logic.

### Outputs

The recorded cumulative motion changes are first converted into a motion heatmap image using a user-selected colormap and then blended with the original video frame via a weighted sum based on the specified weighting parameters, and saved to a temporary file. The retrieved top three peak intensity locations are marked on the blended motion heatmap image with circle, square, and triangle symbols using OpenCV. The base64 module is used to encode the blended motion heatmap image into a base64 format for embedding and display in the motion heatmap image tab in the software. A series of motion changes from the processed frames is converted into a motion heatmap video using the imageio package, and the resulting video is stored as a temporary file on the local disk. The Flask framework is then used to stream and serve the motion heatmap video for display in the motion heatmap video tab in the software, allowing users to playback and monitor the motion changes over time. In addition to these visualization outputs described above, a summary table is created and displayed in the Dash GUI using the Pandas package (The pandas development team, 2020) to summarize the top three peak intensity locations and the total and within-quadrant percentage of regions used, quantifying the spatial distribution and intensity of the animals’ motion. The user has the option to download all these outputs locally for convenient access. The source code, detailed instructions, and video tutorials for AnimalMotionViz are available online at: https://github.com/uf-aiaos/AnimalMotionViz.

### Data

To assess how output from the software aligned with space use and locomotor behavior, sample video files were selected for generation of motion heatmaps and continuous observation of behavior. The sample video data used in this study featured Holstein heifer calves (28-42 days of age) housed in group pens (5 calves/pen) at the heifer rearing facility of the University of Florida Dairy Unit (Hague, FL). Four sample video clips of 5 min in duration were drawn from 24 h video recordings, collected using digital video cameras (Axis M2026-LE Network Camera, Axis Communications, Lund, Sweden) mounted in the center of the outside wall of the pen, approximately 3 m from the ground, with video recorded at 15 frames per second to a network video recorder (Surveillance Station, Synology Inc., Bellevue, WA). These data were originally collected for a research trial where pens varied in effective space allowance (sections of the pen were blocked off to provide 3.7, 4.6, or 5.6 *m*^2^/calf) as described by Marin et al. (2024), representing a range of common on-farm practices for raising dairy calves. For each video file, one observer (JBK) recorded location (within each quadrant), lying, stationary standing, and movement (walking or running) for each calf in the pen using Behavior Observation Research Interactive Software (BORIS; Friard and Gamba, 2016). These data were then summarized at the pen level as the average duration of each behavior (*s*/calf) by quadrant. Quadrants within each pen were then ranked, based on total duration of time per quadrant and duration of time lying, standing, and moving per quadrant.

## Results

The GUI of the AnimalMotionViz application is shown in Figure 2. The software is divided into two main columns: the input section on the left and the output section on the right. The left column allows the user to upload a video and adjust various settings, including video processing parameters and color maps. The right column contains two tabs and a summary table. The motion heatmap image tab displays the final motion heatmap image with the top three peak intensity locations highlighted with colored circle, square, and triangle symbols. The motion heatmap video tab displays the processed motion heatmap video. The user can seamlessly toggle between the tabs. The summary table returns the motion metrics that quantify the motion changes over time. The first three rows detail the colors and symbols used to highlight the top three peak intensity locations, allowing for quick identification and interpretation of the most active areas in the video. These peak locations are important for pinpointing areas of significant activity. The remaining rows show the percentage of the region used in the entire view and within each of the four quadrants. The overall percentage quantifies how much space was occupied by animal movement, providing a baseline measure of overall activity levels. The quadrant-specific percentages are particularly useful for assessing the distribution of movement across different areas of the view, providing insight into spatial preferences or constraints in animal movement. By breaking down the analysis into quadrants, users can identify localized patterns of activity and compare the intensity of movement in different sections of the video.

**Figure 2.**
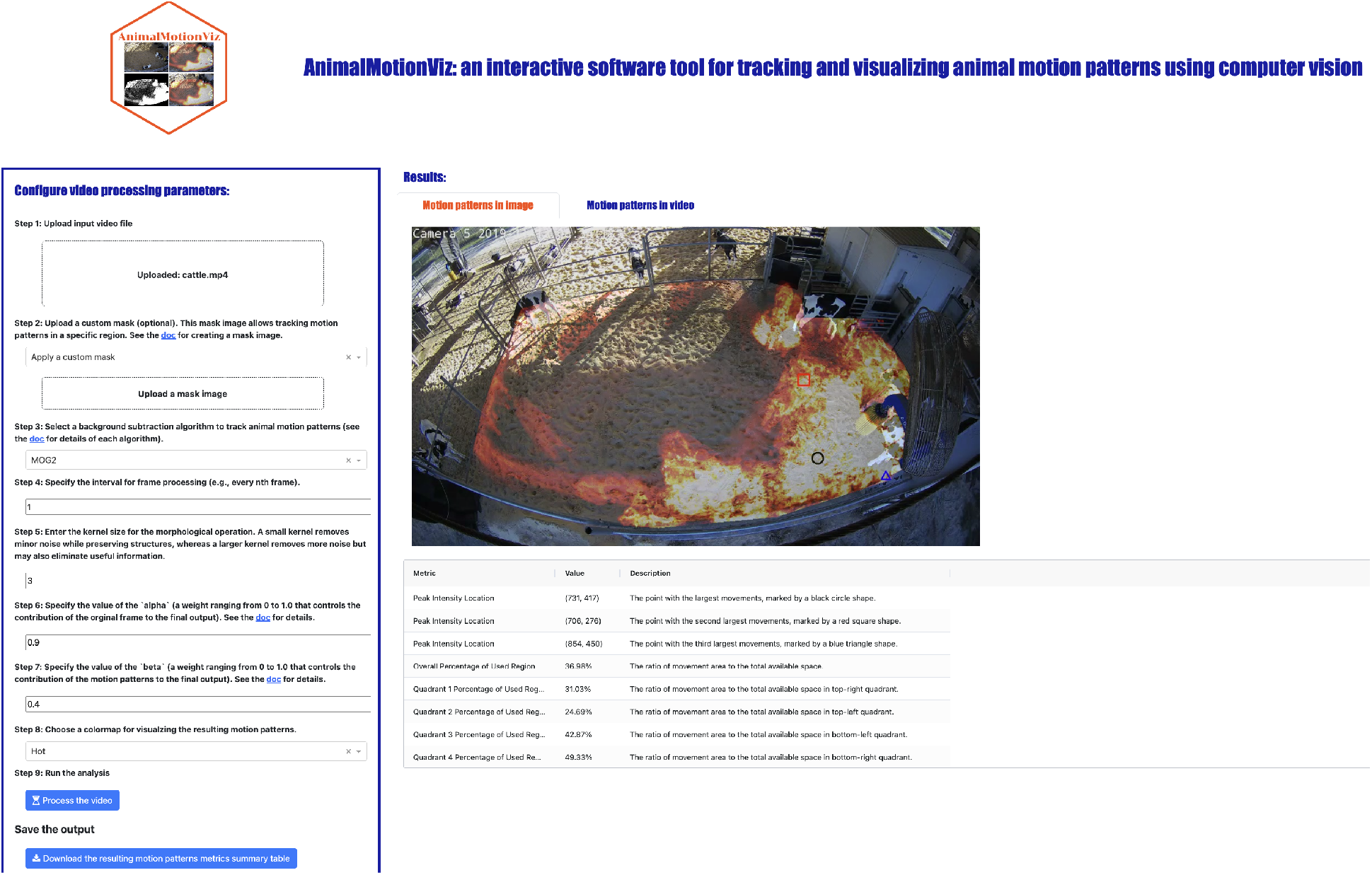
AnimalMotionViz graphical user interface. The left input panel allows the user to upload the input video and configure various parameters. The user also has the option to download the results. The right output panel displays the final motion heatmap image and video, as well as a table with information on peak intensity locations and the percentage of the region used in the entire frame and in each quadrant.

The motion heatmaps of four videos generated by the software are shown in Figure 3. In Figure 3A, the top three peak intensity locations were identified in the upper left quadrant, which aligned with the quadrant with the greatest duration of calf use according to observation (103.4 *s*/calf compared to 3.8 *s*/calf, 102.4 *s*/calf, and 40.7 *s*/calf in the upper right, lower right, and lower left quadrants, respectively), as well as the greatest duration of active movement, such as walking/running (21.7 *s*/calf compared to 2.1 *s*/calf, 6.7 *s*/calf, and 8.0 *s*/calf in the upper right, lower right, and lower left quadrants, respectively). Figure 3B shows the top three peak intensity locations in the upper right quadrant, which was also consistent with the quadrant of the greatest duration of calf use (188.6 *s*/calf compared to 39.6 *s*/calf, 2.3 *s*/calf, and 19.6 *s*/calf in the lower right, lower left, and upper left quadrants, respectively) and the greatest duration of active movement of walking/running (45.9 *s*/calf compared to 5.8 *s*/calf, 9.5 *s*/calf, and 1.7 *s*/calf in the lower right, lower left, and upper left quadrants, respectively). In Figure 3C, the top three peak intensity locations were detected in the upper left quadrant, which corresponded to both the observed longest duration of calf use (162.6 *s*/calf) and active movement (e.g., walking/running: 28.8 *s*/calf), compared to the durations in the upper right (49.6 *s*/calf and 21.4 *s*/calf), lower right (23.8 *s*/calf and 5.7 *s*/calf), and lower left (14.9 *s*/calf and 2.7 *s*/calf) quadrants. Figure 3D shows the top three peak intensity locations in the upper left quadrant, consistent with both the observed longest duration of calf use (115.3 *s*/calf) and active movement (e.g., walking/running: 11.8 *s*/calf), compared to the upper right (15.2 *s*/calf and 1.3 *s*/calf), lower right (114.6 *s*/calf and 7.5 *s*/calf), and lower left (3.9 *s*/calf and 0.8 *s*/calf) quadrants.

**Figure 3.**
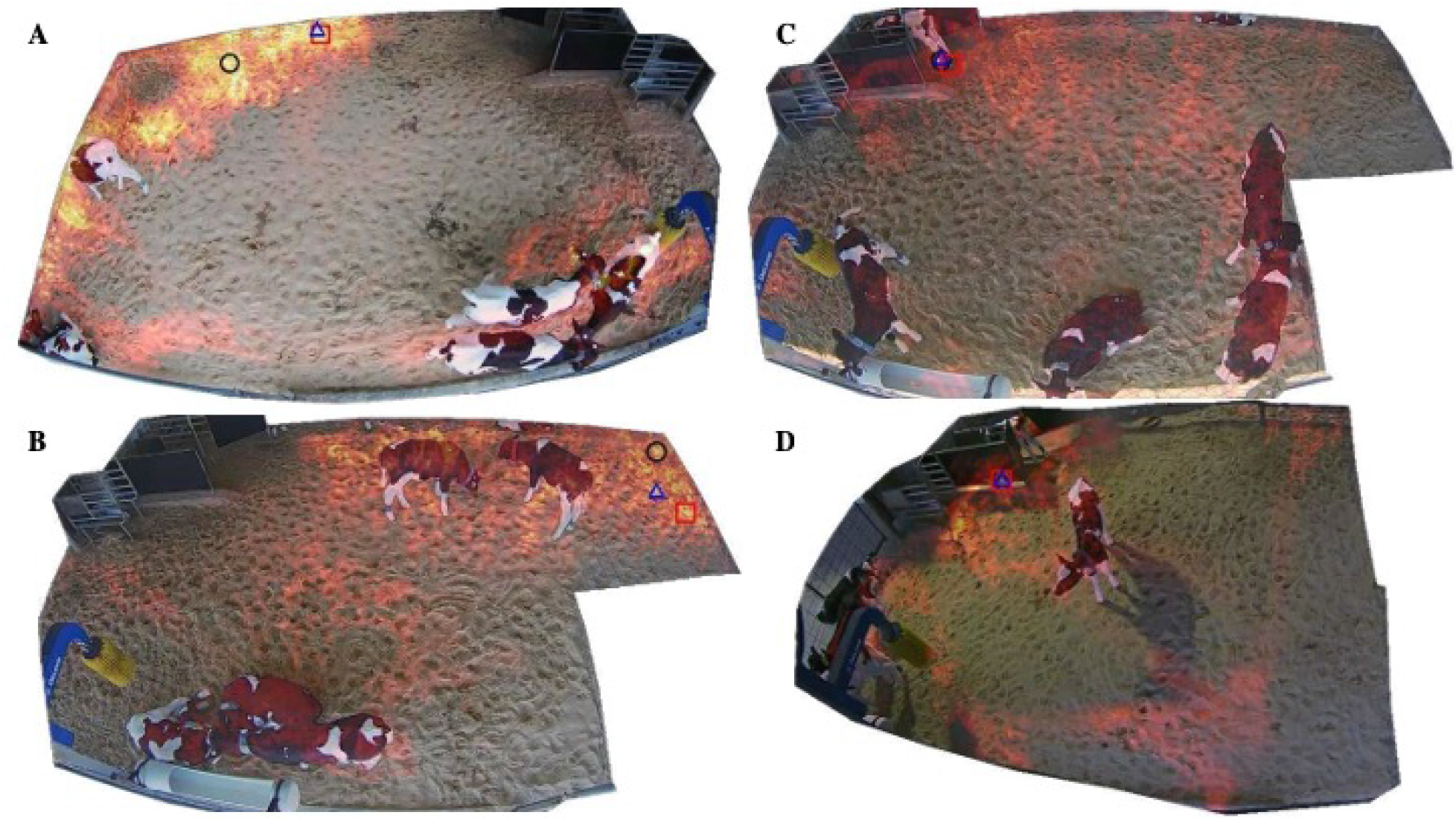
Motion heatmaps tracking the use of pen space for three different group pens (A: 5.6 *m*^2^/calf, B and C: 4.6 *m*^2^/calf, and D: 3.7 *m*^2^/calf). Areas with more motion are shown in warmer colors (gold for the highest motion), while areas with less motion are represented in cooler colors (beige for no motion detected). The top three peak intensity locations are highlighted with black circles, red squares, and blue triangles.

## Conclusion

In the current study, an interactive tool, AnimalMotionViz, was developed to track and visualize animal motion patterns in a video using computer vision. The flexible input column, which can be navigated by mouse clicks, is used to upload a video with no size limitation and to customize the motion pattern tracking parameters. The output column returns a motion heatmap annotated image and video, along with a summary table that uses multiple metrics to quantify the motion patterns relevant to the motion heatmap. The software can be used to understand pen space utilization and space occupation by animals in a barn. The output of the software can enhance management practices and lead to improved pen designs or housing conditions. It also provides an opportunity to study the relationship between space allocation and animal behavior. We believe the development of AnimalMotionViz will accelerate the broader adoption of computer vision systems to further support research developments in precision livestock farming.

## Nonstandard abbreviations used

CNT: CouNT
GMG: Geometric Multi-Grid
GSOC: Google Summer of Code
GUI: graphical user interface
KNN: K-Nearest Neighbors
LSBP: Local SVD Binary Pattern
MOG2: Improved Mixture of Gaussians

## Acknowledgments

This work was supported by the University of Florida startup funds to H.Y.

Sample video files used in this study were collected as part of previous research conducted under IACUC #201909416, as described by Marin et al. (2024).

The authors have not stated any conflicts of interest.

